# A mosaic of independent innovations involving *eyes shut* are critical for the evolutionary transition from closed to open rhabdoms

**DOI:** 10.1101/334532

**Authors:** Simpla Mahato, Jing Nie, David C. Plachetzki, Andrew C. Zelhof

## Abstract

A fundamental question in evolutionary biology is how developmental processes are modified to produce morphological innovations while abiding by functional constraints. Here we address this question by investigating the cellular mechanism responsible for the transition between fused and open rhabdoms in ommatidia of apposition compound eyes; a critical step required for the development of visual systems based on neural superposition. Utilizing *Drosophila* and *Tribolium* as representatives of fused and open rhabdom morphology respectively, we identified three changes required for this innovation to occur. First, the expression pattern of the extracellular matrix protein Eyes Shut (EYS) was co-opted and expanded from mechanosensory neurons to the photoreceptor cells in taxa with open rhabdoms. Second, EYS homologs in these taxa obtained a novel extension of the amino terminus leading to the internalization of a cleaved signal sequence. This amino terminus extension does not interfere with cleavage or function in mechanosensory neurons, but it does permit specific targeting of the EYS protein to the apical photoreceptor membrane. Finally, a specific interaction evolved between EYS and a subset of Prominin homologs that is required for the development of open, but not fused, rhabdoms. Together, our findings portray a case study wherein the evolution of a set of molecular novelties has precipitated the origin of an adaptive photoreceptor cell arrangement.

**Author Summary:** Understanding how adaptive morphologies originate is a central question in evolutionary developmental biology. Once confined largely to arguments about the relative frequencies of protein coding vs. regulatory mutations, numerous studies have since revealed more complex interactions involving alterations in gene expression and novel protein-protein interactions as drivers of novel trait evolution. Our study exploits the genetic amenability of *Drosophila* and utilizes direct comparisons with *Tribolium* to define a set of cellular mechanisms necessary for the evolutionary transition from a fused (*Tribolium*) to an open (*Drosophila*) rhabdom. Our results depict an evolutionary transition involving both non-coding and coding changes that resulted in a novel visual architecture, permitting a subset of diurnal insects to diversify into niches characterized by low light.

## Introduction

Compound eyes are common visual structures found in a wide range of arthropods [1]. Within apposition compound eyes, the ommatidium is the fundamental repeated modular unit required for the detection of light. An ommatidium contains photoreceptor neurons and each photoreceptor has a light gathering organelle known as the rhabdomere. The rhabdom is the collection of all rhabdomeres within a single ommatidium. Rhabdoms may be arranged in two configurations: Open rhabdoms have a pronounced inter-rhabdomeral space (IRS) and are exemplified by the fruit fly *Drosophila melanogaster* (Dm) while fused rhabdoms lack the IRS and are exemplified the flour beetle *Tribolium castaneum* (Tc). In apposition compound eyes, open rhabdoms, coupled with neural superposition, enable an increase in light sensitivity without a commensurate loss of visual acuity [2-4]. Therefore, the evolution of open rhabdoms is thought to have permitted some diurnal dipterans to diversify into niches characterized by low light [5]. Genetic and molecular dissection of *Drosophila* open rhabdoms has identified two key proteins, Eyes Shut (EYS; a.k.a. Spacemaker) and Prominin, that underlie the development of the open rhabdom. EYS and Prominin are elements of a cellular network that generates the IRS involving the secretion of an extracellular matrix, steric hindrance of adhesion and cellular contraction [6-10]. EYS and Prominin are not restricted to open rhabdom species such as *Drosophila* but are instead widely conserved among insects, including species with fused configurations [7, 10], however, the differences in the structure and function of EYS and Prominin that give rise to the evolution of both fused and open rhabdom systems remain unclear.

Here we explore this question by combining functional analyses of EYS and Prominin orthologs from *Drosophila* (Dm) and *Tribolium* (Tc), representatives of open and fused rhabdom systems respectively, and comparative phylogenetic analyses of these loci from selected holometabolous insects. Our findings show that the morphological transformation from a fused to an open rhabdom was based on a mosaic of innovations including: 1) the co-option of rhabdomeric photoreceptor expression from mechanosensory neurons, 2) a change in protein structure of EYS that permitted the targeting of EYS to the apical membrane of photoreceptors, without ablating its function in mechanosensory neurons, and 3) the origination of a novel interaction between EYS and Prominin that facilitated the expansion of the IRS. Moreover, comparative phylogenetic analyses suggest that these molecular innovations may be common among holometabolous insect species with open rhabdoms. Together, our work reveals a set of cellular and molecular innovations required for the adaptive transition from fused to open rhabdoms in compound eye development.

## Results

### The ancestral holometabolous insect possessed a fused rhabdom

Our phylogenetic analyses focus on a selection of phylogenetically informative holometabolous insect species that that represent both fused and open rhabdom configurations. We estimated a well resolved species tree for these taxa using standard methods. We analyzed the evolutionary history of rhabdom configuration among a selection of holometabolous insect taxa using ancestral state reconstruction. Our analyses provide strong support (*P* = 1.0) for the hypothesis that the last common ancestor of holometabolous insects had a fused rhabdom and that transitions of this morphology to the open configuration likely occurred within Diptera on multiple occasions. Therefore, the open rhabdom morphology of *Drosophila* represents an evolutionary innovation (Fig 1).

**Fig. 1.**
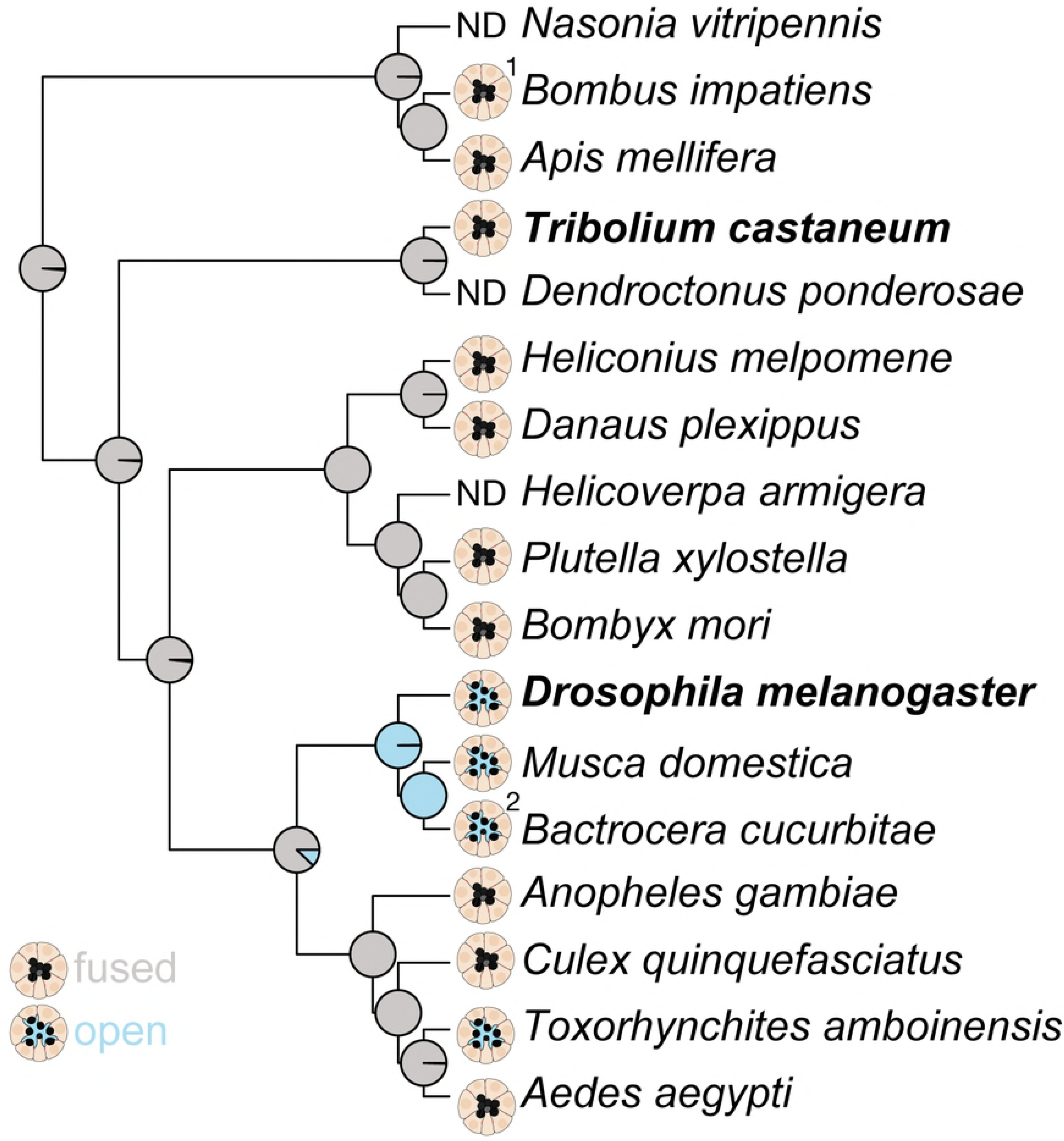
The ancestral holometabolous insect possessed a fused rhabdom. Studies of character evolution using Bayesian simulation mapping show that open rhabdoms have evolved from a fused state at least once in the history of holometabolous insects. The ommatidial arrangement of each taxon is shown. Pie graphs at nodes indicate the posterior probability of reconstructed ancestral states. ND - not determined. ^1^The rhabdom structure was determined for *Bombus dorsalis*. ^2^The rhabdom structure was determined for *Bactrocera dorsalis*.

### Homologous rescue of *eys* mutant phenotypes is dependent upon sensory cell context

Previous data demonstrated that *eys* is expressed in the photoreceptors of open rhabdoms [7, 10] and our genetic dissection of EYS function shows that the loss of Dm-EYS in the open rhabdom of *Drosophila* results in a fused arrangement (Fig 2A,B). We tested the functional equivalence of Dm-EYS and Tc-EYS by assaying their ability to rescue photoreceptor rhabdom arrangement and mechanosensory neuron function in *Drosophila eys* mutants [7, 10, 11]. In *Drosophila*, EYS constitutes part of the extracellular matrix of the IRS that separates each individual rhabdomere within an ommatidium. No rescue of the photoreceptor phenotype was noted upon expression of Tc-EYS in photoreceptors of *Drosophila eys* mutants, in contrast to experiments with Dm-EYS (Fig 2C,D). In Tc-EYS rescue experiments we observed a fused rhabdom and only very limited separation of the photoreceptor stalk membranes juxtaposed to the adherens junctions of photoreceptor cells. In mechanosensory neurons, EYS functions as a protective barrier that safeguards cell shape under environmental stress, heat and changes in osmolarity [11]. As such, *eys* mutants become increasing discoordinated when exposed to high temperatures and this behavior is directly correlated to the loss of mechanosensory function. Upon expression of Tc-EYS in *Drosophila* mechanosensory organs, we found that Tc-EYS was capable of rescuing the mutant mechanosensory defect (S1 Fig), such that the flies were resistant to heat induced discoordination. Further, no morphological defects were observed in mechanosensory organs in Tc-EYS rescue experiments. These results suggest that functional equivalency is dependent upon cellular context; Tc-EYS is apparently fully functional in *Drosophila* mechanosensory neurons but not in rhabdomeric photoreceptors.

**Fig. 2.**
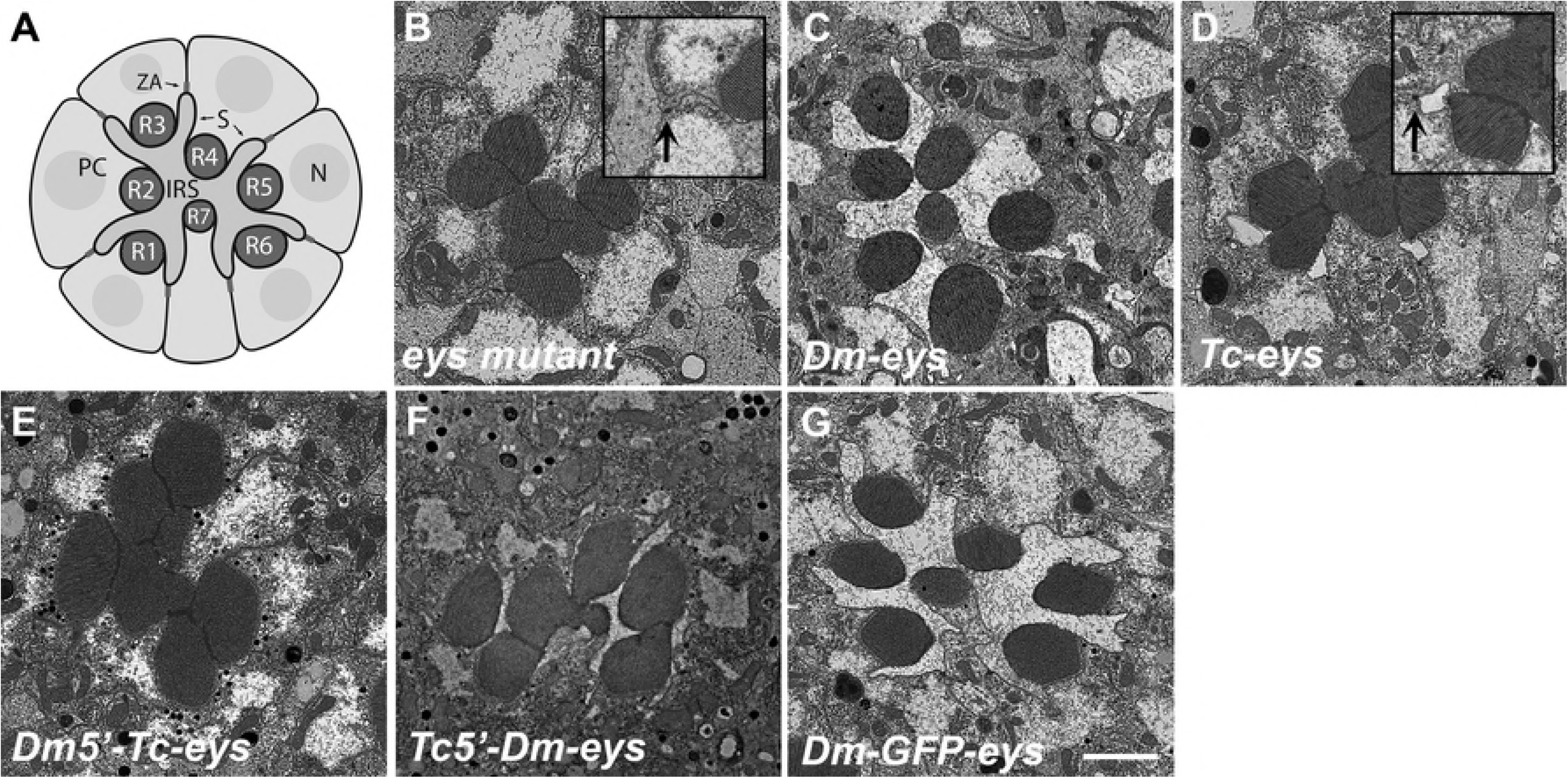
*Drosophila* (Dm) and *Tribolium* (Tc) EYS homologs are not functionally equivalent in photoreceptors. Transmission electron micrographs of adult *Drosophila* ommatidium. A) Schematic of wild type B) *eys* mutant, C) *eys*, Dm-*eys* rescue D) *eys*, Tc-*eys* rescue. E) *eys*, Dm5’Tc-*eys* rescue F) *eys*, Tc5’Dm-*eys* rescue G) *eys*, Dm-GFP-*eys* rescue. Arrows indicate adherens junctions between photoreceptors. IRS – inter-rhabdomeral space. S-stalk membrane. ZA-zonula adherens. PC-photoreceptor cell. N-nucleus. Scale Bar 2 µm.

### Tc-EYS functional rescue correlates with correct spatial localization

We examined the localization of Tc-EYS in both photoreceptor and mechanoreceptor cell types in order to understand the basis for our observation that Tc-EYS could rescue mechanoreceptor, but not photoreceptor, deficiencies in *Drosophila eys* mutants. In photoreceptors, Tc-EYS was detected only in cell bodies, but not at the apical photoreceptor membrane as is Dm-EYS during the critical period of IRS formation (Fig 3A-C). In contrast, in mechanosensory neurons, Tc-EYS localization was identical to that of Dm-EYS (Fig 4A-C). Here, Tc-EYS accumulation was observed at the cavity interface, the junction of the apical surface of the sensory neuron with the lymph space, and inside the lymph space juxtaposed to the ciliary dilation. Thus, the differential ability of Tc-EYS to rescue *eys* mutants directly correlated with its ability to localize correctly in the respective sensory neuron.

**Fig. 3.**
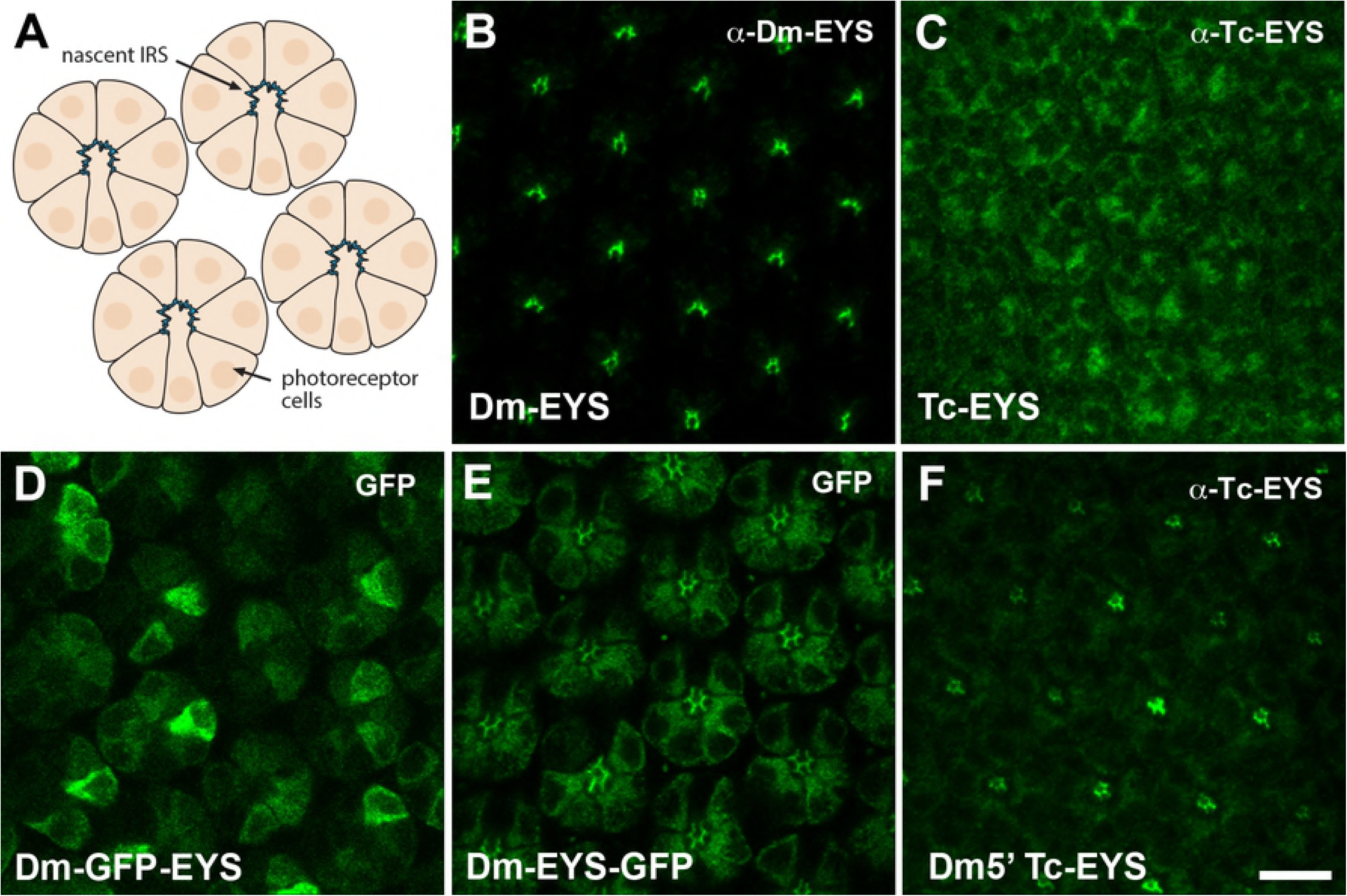
Immunofluorescence of EYS protein localization in 48 hr after puparium formation in wild type photoreceptors. A) Schematic of wild type B) Dm-EYS C) Tc-EYS D) Dm-GFP-EYS E) Dm-EYS-GFP 1 F) Dm5’Tc-EYS. Each panel represents a single confocal slice. Scale Bar 10 µm.

**Fig. 4.**
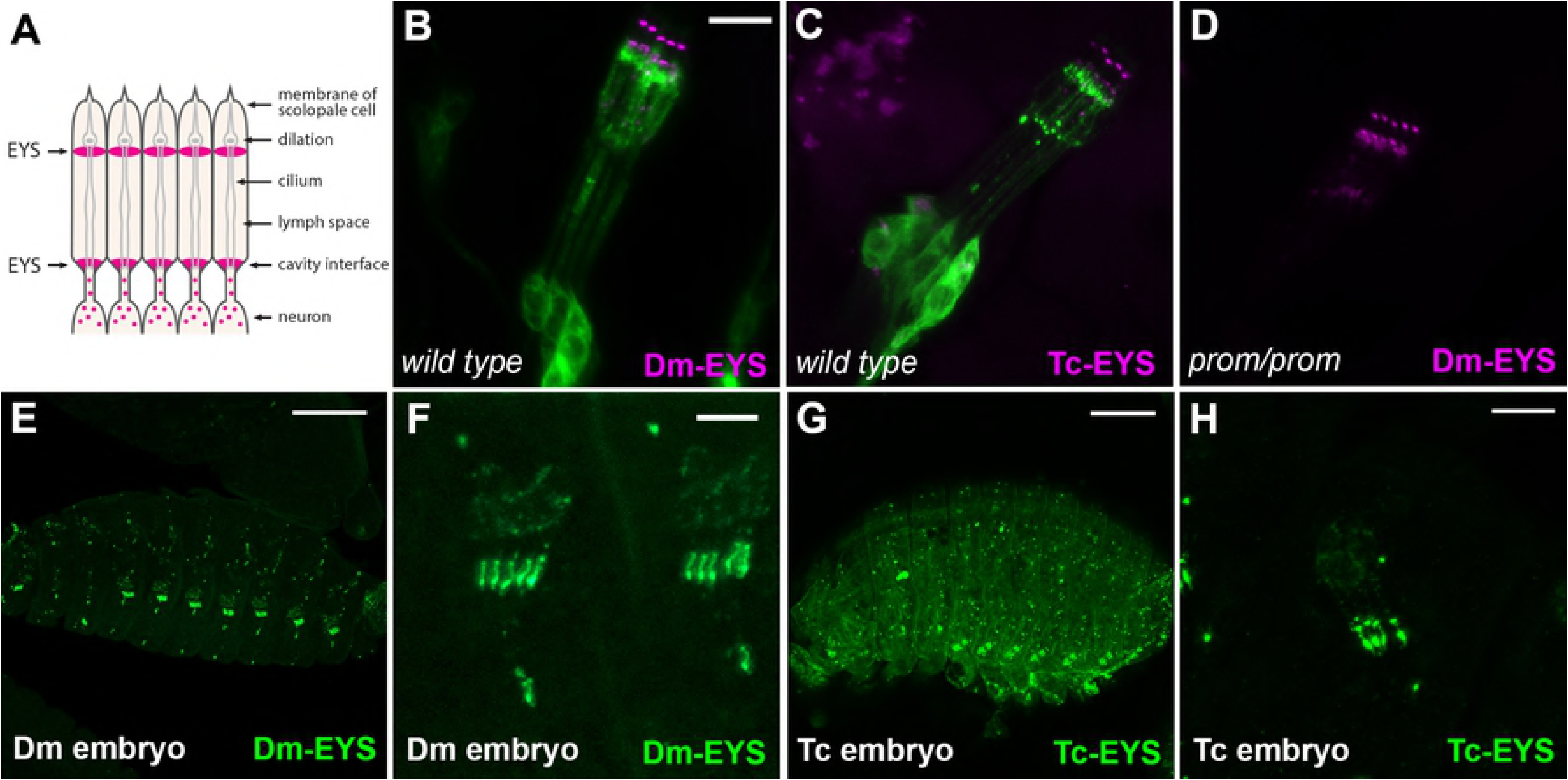
Localization of EYS in mechanosensory neurons, lateral chordotonal organs. A) Schematic of chordotonal structure B) ELAV-GAL4, UAS-CD8-GFP and Dm-EYS (magenta) C) ELAV-GAL4, UAS-CD8-GFP and Tc-EYS (magenta) D) *prominin* mutant and Dm-EYS (magenta) E) *Drosophila* embryo and Dm-EYS (Green) Scale Bar 75 µm. F) *Drosophila* embryo and Dm-EYS (Green), Scale Bar 10 µm. G) *Tribolium* embryo and Tc-EYS (Green), Scale Bar 100 µm. H) *Tribolium* embryo and Tc-EYS (Green) Scale Bar 10 µm. Each panel represents a single confocal slice.

The ability of Tc-EYS to localize and function in both *Tribolium* and *Drosophila* mechanosensory neurons suggested that EYS was co-opted for a novel function in photoreceptors with open configurations. Therefore, we expected to observe Tc-EYS expression in mechanosensory organs of *Tribolium* but not in the retina as suggested by our previous analysis of Tc-EYS mRNA expression [10]. Utilizing our Tc-EYS antibody we find that Tc-EYS is expressed in many neuronal cell types of the *Tribolium* embryonic peripheral nervous system (Fig 4G). In particular, we observed a repetitive pattern in each of the abdominal segments resembling the lateral pentascolopidial chordotonal organs of *Drosophila*. Moreover, cellular localization of Tc-EYS in these cells is highly reminiscent of a similar developmental stage in *Drosophila* embryos (Fig 4E-H). Tc-EYS expression was not detected in *Tribolium* photoreceptors during photoreceptor morphogenesis. Together, these results suggest that the transition from a fused to open rhabdom requires an expanded expression pattern of EYS to photoreceptors.

### A novel amino terminus extension is sufficient for EYS localization to the apical photoreceptor membrane

Whereas presently unknown regulatory changes are implicated in the expanded expression pattern of EYS to photoreceptors, these changes cannot fully account for the adaptation of fused to open rhabdoms because Tc-EYS does not rescue *eys* mutants when expressed in *Drosophila* photoreceptors. Therefore, we reexamined the various structural features of both EYS homologs to determine potential differences that could account for the lack of functional equivalency in photoreceptors. Both Dm-EYS and Tc-EYS contained the same number and arrangement of EGF and Laminin G repeats (Fig 5). Furthermore, as expected for a secreted protein, Tc-EYS contains a typical signal sequence, on the N-terminus just prior to the first EGF domain. In contrast, the N-terminus of Dm-EYS begins with a large stretch of amino acids that appears to internalize a putative signal sequence, which is placed just before the first EGF repeat (Fig 5). This novel extension of the N-terminus was detected across dipteran species with open rhabdoms (Fig 6). In addition, such amino terminal extensions show little sequence similarity and appear to be independently derived.

**Fig. 5.**
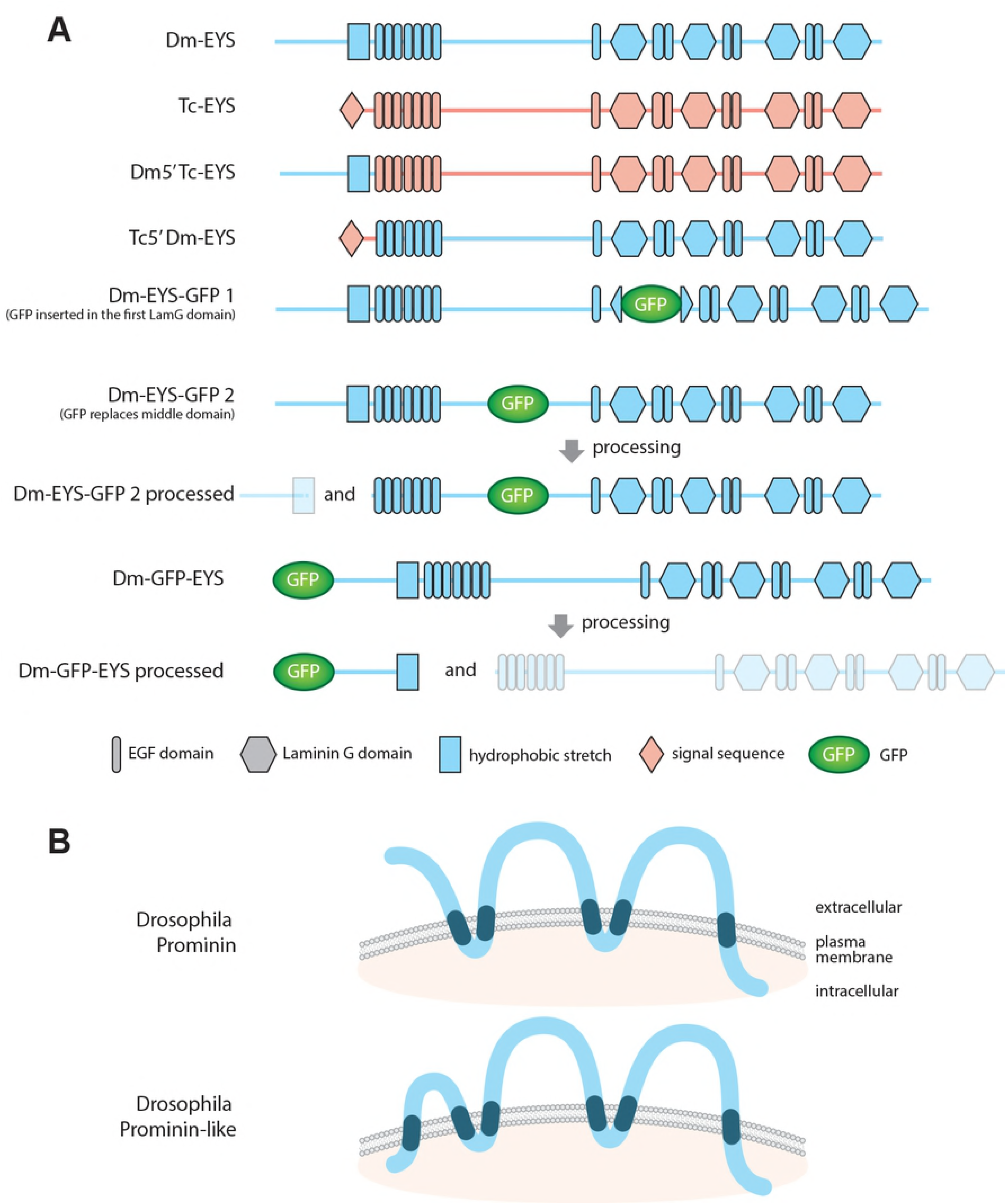
Schematic and Domain Organization of EYS and Prominin proteins.

**Fig. 6.**
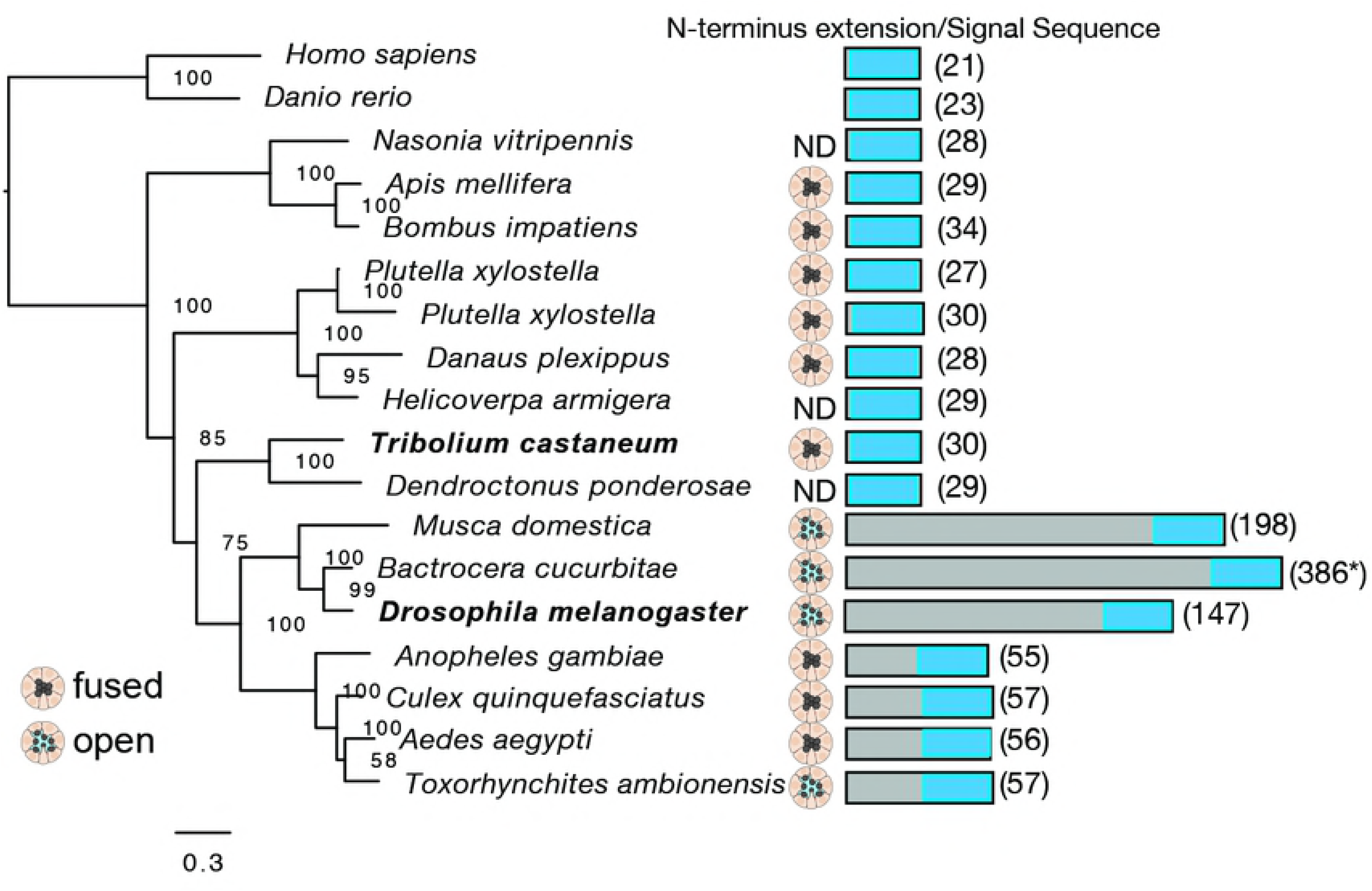
Comparison of EYS amino terminus extension among insects. The amino terminus is defined as the leading amino acids prior to the first cysteine contained within the first EGF domain of the EYS homologs. The blue box represents relative position of predicted signal sequence or internal cleavage site. ND - not determined. * - The extension for *Bactocera cucurbitae* was not experimentally confirmed as compared for *Musca* and *Drosophila*.

To test if this structural variation is necessary and sufficient for EYS function in rhabdomeric photoreceptors, we generated two chimera proteins in which the amino terminus of Tc-EYS and Dm-EYS prior to the first EGF domain were exchanged (Fig 5) and tested for function in *Drosophila* photoreceptors. Replacing the first 30 amino acids of Tc-EYS with the first 147 amino acids of Dm-EYS, prior to the first EGF repeat, resulted in the redirection of Tc-EYS to the photoreceptor apical region during the initial period of IRS formation in wild type photoreceptors (Fig 3F). However, we did not observe any accumulation of the chimera protein in the IRS 24 hrs later (data not shown) and the chimera protein was not capable of rescuing of the *eys* mutant phenotype (Fig 2E). When the *Drosophila* amino terminus was replaced with the amino terminus of *Tribolium*, some of the chimeric protein accumulated in the IRS but there was still a failure of complete separation of the rhabdomeres (Fig 2F). Moreover, immunofluorescence examination demonstrated that this chimera protein, unlike wild type EYS, accumulated abnormally and unevenly throughout the length of the photoreceptor cells (S2 Fig). We conclude that the N-terminus extension is a key feature involved in trafficking EYS to the apical membrane.

### *Drosophila* EYS amino terminus is cleaved upon extracellular release

Secreted EYS is a major component of the IRS. Signal sequences, such as that internalized by a N-terminus extension in Dm-EYS, may be sites for cleavage or could potentially serve as a transmembrane domain [12]. We hypothesized that the internalized signal sequence present in Dm-EYS is a site for cleavage, and the N-terminus extension is removed before subsequent release into the IRS. We first compared the *in vivo* localization of two forms of EYS in which GFP was inserted either before the putative cleavage site, cytoplasmic (GFP-EYS), or after, extracellular (EYS-GFP) [13] (Fig 5). When GFP was located in the putative extracellular region (EYS-GFP) GFP could be detected in the IRS (Fig 3E) and to a lesser extent on the photoreceptor basal lateral membranes. In contrast, utilizing the putative cytoplasmic GFP version (GFP-EYS) GFP was only detected in the photoreceptor cell body and to a lesser extent on the photoreceptor basal lateral membranes. GFP was absent from the IRS (Fig 3D). Nonetheless, the putative cytoplasmic GFP tag of the protein, GFP-EYS, retained the capacity to rescue *eys* mutants (Fig 2G), indicating that the localization pattern was not an artifact of the chimera protein. These data suggested that Dm-EYS N-terminus is cleaved and remains cytoplasmic, while the remainder of the protein is released extracellularly to form the IRS.

To further assess the dynamics of cleavage and secretion of EYS, we leveraged the previous observation that EYS is only detected extracellularly on *Drosophila* tissue culture cells in the presence of Prominin [10]. Neither Prominin nor EYS is normally expressed in *Drosophila* S2 cells [14, 15]. Using our GFP tagged EYS proteins (GFP-EYS and EYS-GFP), we first tested their respective capacities to be secreted and accumulate on S2 cells when co-transfected with Dm-Prominin. In both cases, we observed the accumulation of EYS, as detected by EYS immunofluorescence, on the extracellular region of the cell (Fig 7B,E). The known epitope/antigen for the EYS antibody, mAb21A6, has been mapped to the putative extracellular region of EYS between the fifth and seventh EGF repeats of the amino terminus [10]. However, there was a defined difference with respect to the localization of GFP. Only when the GFP tag was located in the putative extracellular region (EYS-GFP) did we observe accumulation of GFP extracellularly and subsequent colocalization with EYS immunofluorescence (Fig 7A-C). There was no indication that the putative intracellular GFP tag of GFP-EYS was secreted or released extracellularly (Fig 7D-F). Furthermore, when we examined the intracellular pool of each tagged protein, we observed two distinct signals for GFP and EYS with the GFP-EYS construct, indicating that GFP is removed from the GFP-EYS chimera protein. Conversely, we observed only a single colocalization signal with EYS-GFP (S3 Fig). The cleavage was also verified by Western analysis. In cell extracts expressing GFP-EYS we detected a proteolytic cleavage product associated with GFP-EYS of the expected size of GFP plus the additional 147 amino acids prior to and adjacent to the hydrophobic stretch (Fig 8 and Fig 5). Together, our results demonstrate that the internalized signal sequence in Dm-EYS is a cleavage site which is processed before EYS can be secreted. In addition, Prominin is required for the retention of secreted EYS on the extracellular surface, but the processing of EYS occurred in the absence of Prominin in cell culture (Fig 8). This finding is consistent with our observation that the loss of Prominin had no effect on the localization of Dm-EYS in mechanosensory organs (Fig 4D) or the ability of Dm-EYS to be targeted to and secreted in the absence of Prominin [10].

**Fig. 7.**
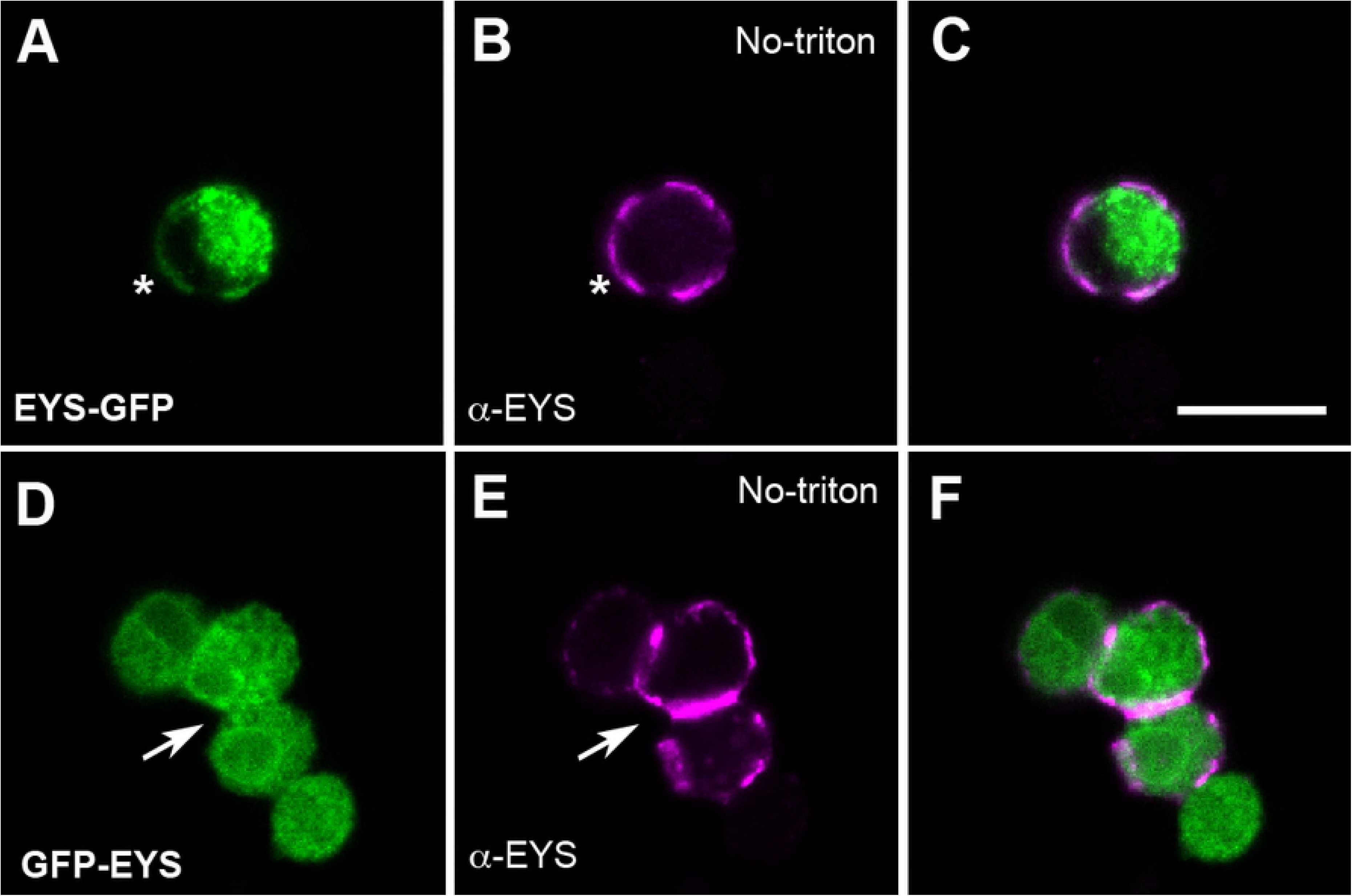
Differential localization of GFP, of GFP epitope tagged versions of Dm-EYS in non-permeabilized tissue culture cells. *Drosophila* S2 cells co-transfected with either EYS-GFP (A-C) or GFP-EYS (D-F) with Dm-Prominin. GFP localization (A,D) was compared to EYS detection with an antibody (B,E) in the absence of detergent, no cell permeabilization. Asterisk indicates colocalization of GFP and EYS epitope. Arrow indicates the accumulation of EYS in the absence of corresponding accumulation of GFP. Scale Bar 10 µm.

**Fig. 8.**
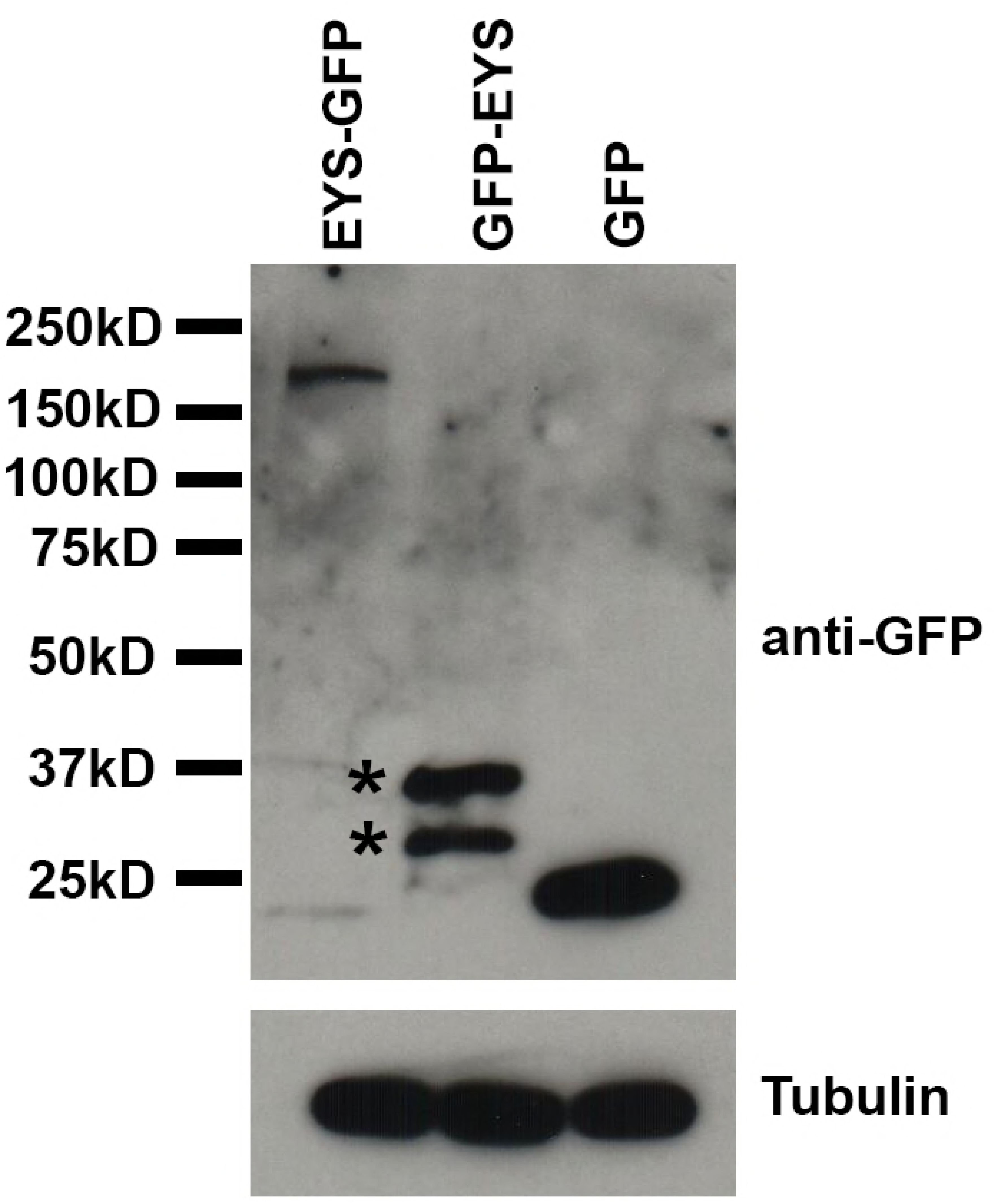
Dm-EYS proteolytic cleavage products are detected in *Drosophila* tissue culture cells. Immunoprecipitation and detection of GFP from *Drosophila* S2 cells transfected with either EYS-GFP, GFP-EYS, or GFP in the absence of Dm-Prominin. Asterisks indicate intracellular GFP containing cleavage products associated only with GFP-EYS.

### IRS formation is dependent upon species specific interactions and a subset of Prominin paralogs

The failure of Tc-EYS to rescue the *eys* photoreceptor mutant even when properly directed to the apical membrane suggested that additional mechanisms are required for the transition from a fused to an open rhabdom. Previous data has demonstrated that the interaction between EYS and Prominin is critical for IRS formation [8, 10]. In particular, Prominin was proposed to be a “receptor” for EYS ensuring proper distribution of EYS during the transformation of the photoreceptor apical membrane into a rhabdomere. Therefore, the lack of rescue observed with Tc-EYS or Dm5’Tc-EYS could be due to a failure of Tc-EYS to interact with *Drosophila* Prominin. To test this possibility, we tested whether either Tc-EYS or Dm5’Tc-EYS could be detected extracellularly on S2 cells in the presence of Dm-Prominin. In both cases, even though protein expression can be detected in S2 cells, neither protein was detected on the surface of the cell, supporting the idea that the interaction between EYS and Prominin required for the proper secretion and positioning of EYS may be highly specific (S4 Fig).

Given the specific interaction between Dm-EYS and Dm-Prominin, we further tested whether open rhabdoms may also correlate with structural changes in Prominin. We performed genome searches and phylogenetic analyses of Prominin homologs from selected holometabolous insects. Our analysis revealed two distinct clades of Prominin homologs across holometabolous insects, Prominin and Prominin-like, as known from *Drosophila* (S5 Fig). Prominin-like loci were lost in a few lineages, but Prominin was present in each species, with duplications noted in hymenopteran species. Thus, the presence of Prominin or Prominin-like did not correlate with the presence of open or fused rhabdoms. Structurally, the analyses of Prominin and Prominin-like proteins demonstrate well supported, highly divergent transmembrane domain structures. Whereas Prominin clade proteins contain five transmembrane domains, Prominin-like has a predicted sixth transmembrane domain. Given the presence of structural differences between Prominin orthologs, we tested whether Prominin and Prominin-like are functionally equivalent. Utilizing similar expression conditions, Dm-Prominin-like was not capable of rescuing prominin mutants (Fig 9A-C), even though Dm-Prominin-like localized to the apical photoreceptor membrane during rhabdomere morphogenesis (S6 Fig). Together, these results reveal structural diversity and a lack of functional equivalency among Prominin paralogs. Interestingly, this pattern of structural variation among holometabolous insect Prominin homologs suggests the potential for novel, species specific interactions between EYS and Prominin in the adaptive evolution of rhabdom architectures.

**Fig. 9.**
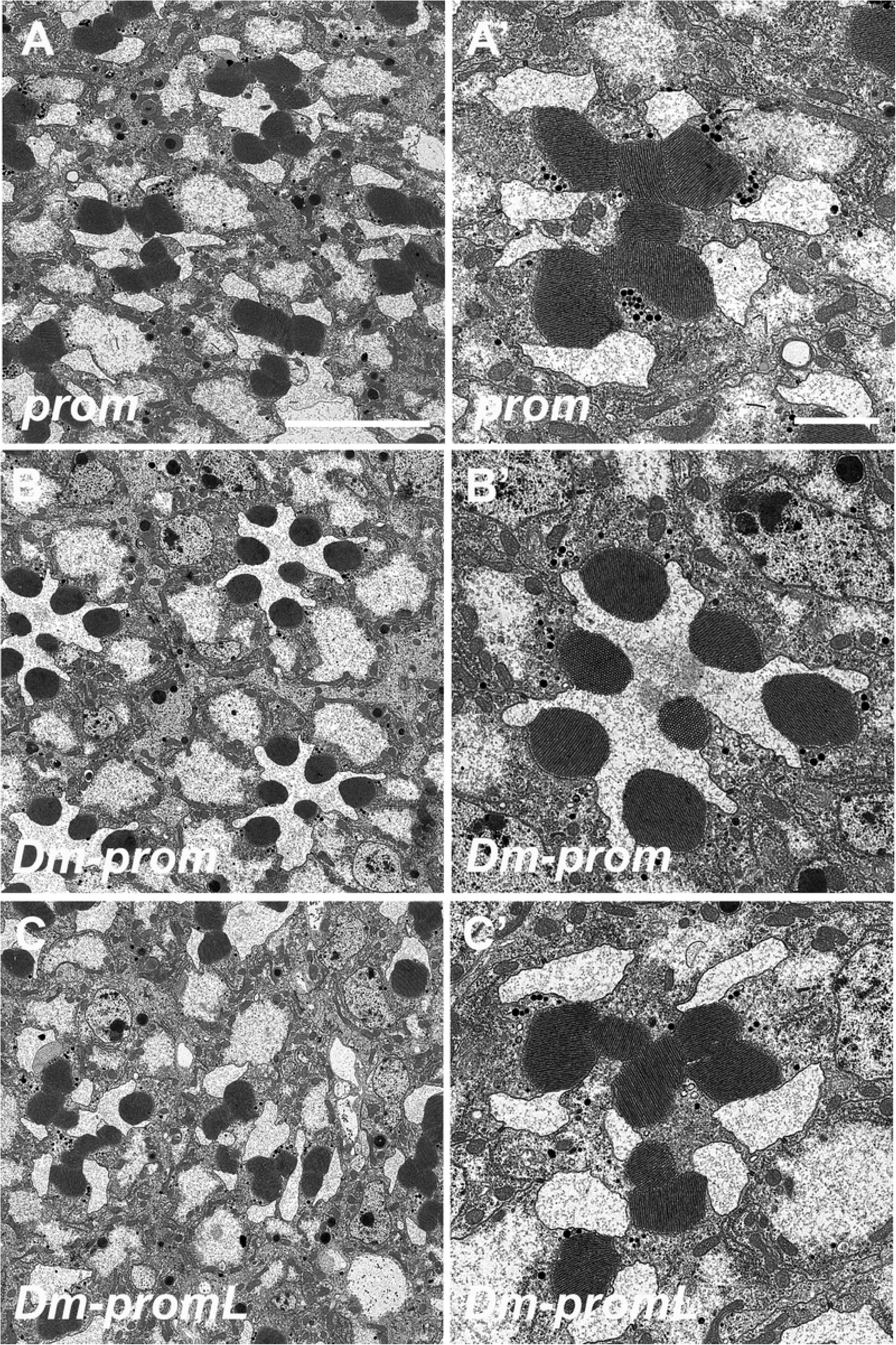
Prominin and Prominin-like are not functionally equivalent. Transmission electron micrographs of adult *Drosophila* ommatidia. A) *prom* mutant B) Rescue with Dm-prominin C) Rescue with Dm-prominin-like. Scale Bars 5 µm and 2 µm.

## Discussion

### Open rhabdoms are evolutionary innovations based on the co-option of EYS from ancestral roles in mechanosensory neurons

Phylogenetic comparative analyses strongly support the hypothesis that the open rhabdom configuration is a derived condition in holometabolous insects (Fig 1). Our data indicate that the role of EYS in mechanosensory neurons is conserved between *Drosophila* and *Tribolium* and was co-opted in *Drosophila* photoreceptor cells for the generation of the IRS. We found that EYS expression and subcellular localization is conserved in mechanosensory neurons of both *Tribolium* and *Drosophila*, in particular the lateral chordotonal organs in developing embryos. Interestingly, despite the differences in protein sequences, proteins from either species were capable of functioning in mechanosensory neurons, but only the *Drosophila* protein (Dm-EYS) was functional in photoreceptors.

EYS is an evolutionary conserved molecule found outside of insects. EYS homologs are characterized by an amino terminus containing EGF repeats, a non-conserved central domain, and a carboxy terminus of alternating EGF and LaminG domains. However, only in Diptera is EYS characterized by an amino terminus extension that leads to the internalization of the signal sequence. Like *Tribolium*, both human and Zebrafish EYS have typical, exposed, signal sequences of 21 and 23 amino acids, respectively, which function in secretion [16-19]. Moreover, the common feature of all EYS homologs studied to date is expression in ciliated neurons. In deuterostomes, EYS expression is limited to ciliary photoreceptor neurons where it is involved in maintaining the integrity of the ciliary photoreceptors [16-20]. In insects, mechanosensitive neurons are also ciliated and EYS functions to preserve chordotonal neuron cell shape in the face of temperature, osmotic or chemical stress [11]. In both systems, EYS is involved in protecting the integrity of ciliary projections among different types of sensory neurons. Through co-option for function in rhabdomeric photoreceptors, EYS in open rhabdom systems took on a new role as a major component of the IRS.

### Internalization of signal sequence for cleavage: a novel change for the adaptation from fused to open rhabdom

Our results also show that the evolution of a novel N-terminus extension that effectively buries the secretion signal sequence in *Drosophila* EYS was also essential for the transition to an open rhabdom configuration. Signal sequences for export are typically 15-30 amino acids in length, have low sequence homology, but contain three identifiable domains: a basic, hydrophobic, and polar domain. In addition to signaling a given protein for secretion, signal sequences can provide other forms of information, including trafficking [12, 21]. In *Drosophila*, we demonstrated that an alteration in the localization of the EYS signal sequence is critical for the transition from fused to open rhabdoms. Across holometabolous insects EYS has a typical 15-30 amino acid signal sequence at the N-terminus of the protein. In Dm-EYS and other dipterans, EYS homologs are characterized by the internalization of this signal sequence due to a novel N-terminus extension that can be over 100 amino acids in length. Our data show that this N-terminus extension does not ablate the signal sequence, but it is responsible for proper targeting of EYS to the apical membrane and is cleaved prior to secretion. This N-terminus extension has no effect on function in mechanosensory neurons but is necessary for proper targeting of EYS to the apical membrane in *Drosophila* photoreceptors. When Tc-EYS is expressed in *eys* mutant photoreceptor cells, only a slight separation of the rhabdomeric stalk membranes closest to the adherens junctions (Fig 2D) is observed. Therefore, the signal sequence alone was able to direct a limited portion of Tc-EYS to be secreted, while the majority of Tc-EYS remained intracellular in regions other than the apical membrane at the critical period of IRS formation (Fig 3C). The replacement and addition of the N-terminus of Dm-EYS to Tc-EYS resulted in the chimeric protein being targeted to the photoreceptor apical surface. However, despite correct targeting, Dm5-Tc-EYS was not capable of rescuing *eys* mutants, suggesting other factors are required. Conversely, the replacement of the Dm-EYS N-terminus with the Tc-EYS signal sequence also resulted in an incomplete separation, but in these experiments the failure to rescue *eys* mutants was due to some mis-targeting of EYS to other regions of the developing photoreceptors (S2 Fig).

A typical signal sequence is not sufficient to target EYS to the apical photoreceptor membrane, suggesting a novel pathway for trafficking EYS to the apical surface. Moreover, this putative pathway may not be limited to targeting EYS in open rhabdom systems. A second feature of open rhabdoms is that the apical photoreceptor membrane is divided into two distinct domains, the rhabdomere and the stalk membrane, the latter of which is devoid of microvilli. The generation of the stalk membrane is dependent on the transmembrane protein Crumbs [22-24], which localizes to the entire stalk membrane. Like Dm-EYS, Dm-Crumbs has an internalized cleavable signal sequence with an 83 N-terminus extension. This extension, like EYS, is not present in Crumbs homologs from species with fused rhabdoms. Crumbs proteins with fused rhabdoms including *Tribolium castaneum, Aedes aegypti* and *Nasonia vitripennis* all have typical signal sequences of 22, 25, and 22, amino acids, respectively, as also seen with human Crumbs proteins [25]. To date the sufficiency and necessity of this N-terminus extension in Crumbs has not been tested in *Drosophila* photoreceptors.

### Prominin orthologs are not interchangeable

It is intriguing that the chimeric protein Dm5-Tc-EYS (Tc-EYS with the N-terminus extension of Dm-EYS) localized correctly to the apical membrane in *Drosophila* photoreceptors but was still unable to rescue the *eys* mutant as it failed to generate an inter-rhabdomeral space. Conversely, the chimera Tc5-Dm-EYS, which lacks the targeting signal altogether but has an otherwise wild type EYS, was capable of generating a partial IRS. Why is correct targeting to the apical membrane not sufficient to rescue *eys* mutants? Prominin has been hypothesized to be a receptor for EYS, but detailed studies of Prominin function, or how it promotes the secretion and/or accumulation of EYS is not known. We hypothesize that, together with the change in tissue expression and protein structure in Dm-EYS, an additional specific protein-protein interaction between EYS and Prominin must have evolved to facilitate the proper development of the IRS in the open rhabdom system in *Drosophila*. Our results from cell culture and *in vivo* studies show that Prominin orthologs examined are not functionally equivalent and we confirmed the species-specific nature of EYS-Prominin interactions in cell culture experiments where Tc-EYS and Dm-Prominin did not interact in a manner that resulted in the proper deployment of EYS, but Dm-EYS and Dm-Prominin were able to do so. Our results, also are an agreement with previous experiments that tested the functional equivalency of the human homologs EYS and Prominin-1 [8] in *Drosophila* photoreceptors. Whereas, human Prominin-1 was functionally equivalent to Dm-Prominin, human EYS alone did not rescue *eys* mutants.

### Conclusions

Understanding how adaptive morphologies originate is a central question in evolutionary developmental biology. Once confined largely to arguments about the relative frequencies of protein coding vs. regulatory mutations as drivers of novel trait evolution [26-28], our study adds yet another dimension to this understanding as we outline a mosaic of concomitant mutations that underlie the evolution of an open rhabdom from a closed rhabdom configuration in scotopic *Drosophila*. These alterations, each necessary, profoundly affect the cell biology of developing photoreceptor cells and result in the expansion of the IRS, driven by the proper deployment and secretion of EYS. First, the expression pattern of EYS must itself expand from being limited mostly to mechanosensory organs, as in *Tribolium*, to include photoreceptor expression, as in *Drosophila*. In order for EYS to be targeted to the apical membrane, the evolution of an N-terminus extension was required. This N-terminus extension, present in taxa with open rhabdoms, but absent from taxa with closed configurations, does not directly contribute to the IRS, but is necessary for proper targeting of EYS in the apical membrane. Finally, specific protein-protein interactions evolved between EYS and Prominin that facilitate the proper deployment of secreted, cleaved, EYS to the IRS. Our data indicate an evolutionary transition involving both non-coding and coding changes that resulted in a novel visual architecture permitting a subset of diurnal dipteran lineages to diversify into niches characterized by low light.

## Methods

### Species Tree Estimation

We selected 16 high-quality genome and transcriptome datasets from species representing the major lineages of holometabolous insects where ommatidial morphology was known. We constructed a set of orthologs for phylogenomic analysis using the partitions described in [29] and our custom reciprocal BLAST scripts. Briefly, our procedure first extracted the sequence data from each partition for the reference species *Drosophila melanogaster* and BLASTed back against the dmel-R6 protein set [30]. The resulting top hit was in turn BLASTed against each of our 16 species protein datasets. Resulting best hits were then blasted back against the dmel-R6 protein set and each sequence for which both the original and secondary blast searches hit the same protein in the dmel-R6 protein set was retained as a representative ortholog for that species. Individual partitions that had greater than 90% taxon occupancy were aligned using MAFTT [31] under default parameters, trimmed using gblocks_wrapper.pl and concatenated using custom scripts. The resulting phylogenomic dataset included 37 partitions and had a total length of 16,976 amino acid positions. Phylogenetic reconstruction was done using Phylobayes MPI [32].

### Ancestral State Reconstruction

For ancestral state reconstruction, we encoded each species as having either a fused or an open rhabdom. We then reconstructed the ancestry of the fused/open character state using make.simmap implemented in the R package phytools (https://doi.org/10.1111/j.2041-210X.2011.00169.x). Character state transitions were modeled under equal rates and 10,000 simulated histories were examined.

### Gene Tree Estimation

The same protein sets used for phylogenetic reconstruction were also used for EYS and Prominin gene tree estimation, with the exception that vertebrate model taxa were also included. Here, BLAST [33] searches using EYS homolog (AAZ83988.3) and Prominin (NP_001286843.1), both from D. melanogaster, were conducted and the top 10 hits above an e-value of 0.001 were retained. Sequences were aligned using MAFFT under default parameters and estimated phylogenies using RaXML under the AUTO model setting. Initial phylogenies for both EYS and Prominin included several clades of closely related paralogs (data not shown). From these large gene family trees, we then pruned the focal EYS or Prominin clades from these trees, realigned these sequences in MAFTT under default parameters. Phylogenetic estimation was then done using RaXML with 20 random starts under the AUTO model setting. We also conducted 1000 bootstrap replicates and support was assessed by mapping bootstrap replicates onto the best scoring tree from the previous step. All data and scripts used in the phylogenetic aspects of the study are available at https://github.com/plachetzki/EYS_PROM.

### *Drosophila* stocks and cDNAs

All crosses and staging were performed at 23°C. *Drosophila* stocks used in this study include: UAS-Dm-*eys*, UAS-Dm-*prominin, prom*^1^, *eys*^1^ [10], prom-gal4 [8]. chp-gal4 (BDSC#47686), elav-gal4 (BDSC #458), UAS-mCD8-GFP (BDSC #5137), Dm-EYS-GFP 1 (BDSC # 63162) were obtained from the Bloomington Drosophila Stock Center. The following stocks were created for this study UASattB- Dm5’Tc-*eys*, UASattB-Tc5’Dm-*eys*, UASattB-Tc-*eys*, UASattB-Tc-*prominin-like* and inserted into y-attP-3B (BDSC#24871), UASattB-Dm GFP-EYS, UAS-Dm EYS-GFP 2 and UAS-Dm-*prominin-like*. cDNA representing Tc-*eys* was constructed from RT-PCR reactions (SuperScript III First Strand Synthesis, Invitrogen) from total RNA isolated from *Tribolium v*^*W*^ stock, respectively. cDNA representing isoform Dm-*prominin-like B* was obtained from the Drosophila Genomics Resource Center (DGRC # 130400). The 5’ ends of *Musca domestica* and *Toxorhynchites amboinensis* were confirmed by 5’RACE (Generacer Core Kit, Invitrogen). *Musca domestica* were obtained from Carolina Biologicals (#Item #144410) and mRNA and total RNA for *Toxorhynchites amboinensis* was obtained from Dr. J. Pitts (Baylor University). Chimeric cDNAs were produced utilizing NEBuilder HiFi assembly (New England Biolabs).

### Antibody Production

Tc-eys antibody production: The first 310 amino acids of the ORF was fused to GST and used as an antigen in rats (Cocalico Biologicals). Dm-Prominin-like antibody production: - The C-terminal peptide SERDREHVPLANVPKK (produced in the laboratory of Dr. Charles Zuker) was used as the antigen in rabbits.

### Transmission electron microscopy, immunofluorescence staining, and imaging

*Drosophila* eye samples were prepared for transmission electron microscopy (TEM) as previously described [34]. All crosses were maintained at 23°C and adult heads were fixed within 8h after eclosion. Pupal retinas were staged at 23°C, dissected in PBS, and fixed in PBS containing 4% formaldehyde for 10min. *Drosophila* embryos were collected and fixed as described in [35] *Tribolium* embryos were collected and fixed as described in [36] but late stage embryos were selected and manually dechorionated before incubation with primary antibody. The primary antibodies used were: mouse anti-EYS (mAb 21A6, 1:50, Developmental Studies Hybridoma Bank); rabbit anti-Prom (1:100) [10]; rat anti-Tc-EYS (1:50), Rhodamine (1:200) or Alexa Fluor 647 (1:50) conjugated phalloidin (Life Technologies) was used for the detection of F-actin. The FITC or RX secondary antibodies (1:200) were obtained from Jackson ImmunoResearch Laboratories. Confocal images were taken on a Leica TCS SP5 and TEM was performed with a JOEL 1010 and all images were processed in Adobe Photoshop.

### Cell Transfections, Co-immunoprecipitations and Westerns

Cell transfection assays and immunofluorescence were performed as previously described [8, 10], utilizing Qiagen Effectene. S2 DGRC cells were obtained from the Drosophila Genomics Resource Center (DGRC #6). A GFP nanobody (Allele) was used to immunoprecipitate GFP containing proteins from S2 cell extracts according to the manufacture instructions. Proteins were separated on Mini-Protean TGX gels (BIO-RAD) and transferred to Immobilon membranes (Millipore). Antibodies utilized were rabbit anti-GFP (ab290-Abcam) and mouse monoclonal anti-alpha Tubulin (T9026-Sigma). Signal detection was achieved with use of a HRP-conjugated anti-mouse or anti-rabbit secondary antibody (1:5000) (Jackson ImmunoResearch Laboratories) combined with Superscript West Pico Chemiluminescent Substrate (Thermo Scientific).

### Signal sequence and Transmembrane domain analyses

The following programs were utilized to analyze protein structures: SignalP 4.1 Server, HMMER, and TMHMM Server v. 2.0.

## Acknowledgments

We thank Dr. Jason Pitts for *Toxorhynchites ambionensis* material and sequence data and the Bloomington Drosophila Stock Center and the Drosophila Genomics Resource Center for reagents. We thank Dr. J. Powers and the Indiana University Light Microscopy Center for assistance with image generation.

## Supporting Information Legends

**S1 Fig. The *Tribolium* EYS homolog can functionally rescue *eys* deficient ciliary mechanoreceptor neurons.** Expression of Tc-EYS in neurons (ELAV-GAL4) was capable of rescuing the mechanosensory defect associated with *eys* mutants. Four separate trials of each genotype, males and females, were heat-shocked for 1 hr at 37°C, allowed to recover for 15 minutes and then each vial was banged twice and allowed to recover for five minutes. The animals were scored for the following characteristics: *paralyzed* – on back or side on the floor of vial, *standing* – upright on floor of vial, *climbing* – on the side of the vial. The percent animals and SEM were calculated for each genotype.

**S2 Fig. Mis-localization of Tc5’-DmEYS in developing photoreceptors.** Immunofluorescence of Tc5’ DmEYS protein and localization in 52 hr APF wild type photoreceptors. A-C) Apical section of developing photoreceptors stained for Tc5’-DmEYS (antibody against Dm-EYS) and Phalloidin to mark F-actin. D-F) Lateral view of developing photoreceptors stained for Tc5’-DmEYS (antibody against Dm-EYS) and Phalloidin to mark F-actin. Asterisks mark abnormal accumulation of EYS protein. Each panel represents a single confocal slice. Scale Bar 10 µm.

**S3 Fig. Differential localization of GFP, of GFP epitope tagged versions of Dm-EYS in permeabilized *Drosophila* tissue culture cells.** *Drosophila* S2 cells transfected with either EYS-GFP (A-C) or GFP-EYS (D-F). GFP localization (A,D) was compared to EYS detection with an antibody (B,E) in the presence of detergent, cell permeabilization. Scale Bar 10 µm.

**S4 Fig. Tc-EYS and Dm5’-Tc-EYS do not accumulate extracellularly in the presence of Dm-Prominin in Drosophila tissue culture cells.** *Drosophila* S2 cells transfected with either Tc-EYS (A), Dm5’-Tc-EYS (B), and Tc5’-Dm-EYS in the presence of Dm-Prominin and cytoplasmic GFP. In the absence of detergent (Triton) only Tc5’ Dm-EYS is detected extracellularly on the transfected cells. There is no difference between background fluorescence between transfected and non-transfected cells containing Tc-EYS or Dm5’-Tc-EYS. Scale Bar 10 µm.

**S5 Fig. Prominin Phylogeny among holometabolous insects.** Prominin and Prominin-like proteins are present insects with both open and closed rhabdoms.

**S6 Fig. Localization of Prominin and Prominin-like in *Drosophila* photoreceptors.** A) Immunofluorescence of Dm-Prominin (green) in wild type photoreceptors at 48 hrs after puparium formation countered stained with Dm-EYS (magenta) and F-Actin (blue). B) Immunofluorescence of Dm-Prominin-like (green) in *prominin* mutant photoreceptors at 48 hrs after puparium formation countered stained with Dm-EYS (magenta) and F-Actin (blue). Scale Bar 10 µm.

